# Rational Prediction of PROTAC-compatible Protein-Protein Interfaces by Molecular Docking

**DOI:** 10.1101/2023.02.16.528819

**Authors:** Gilberto P. Pereira, Brian Jiménez-García, Riccardo Pellarin, Guillaume Launay, Sangwook Wu, Juliette Martin, Paulo C. T. Souza

## Abstract

Proteolysis targeting chimeras (PROTACS) are heterobifunctional ligands that mediate the interaction between a protein target and an E3 ligase, resulting in a ternary complex whose interaction with the ubiquitination machinery leads to target degradation. This technology is emerging as an exciting new avenue for therapeutic development, with several PROTACS currently undergoing clinical trials targeting cancer. Here, we describe a general and computationally efficient methodology combining restraint-based docking, energy-based rescoring, and a filter based on minimal solvent-accessible surface distance to produce PROTAC-compatible PPIs suitable for when there is no *a priori* known PROTAC ligand. In a benchmark employing a manually curated dataset of 13 ternary complex crystals, we achieved accuracy of 92% when starting from bound structures, and 77% when starting from unbound structures, respectively. Our method only requires that the ligand-bound structures of the monomeric forms of the E3 ligase and target proteins be given to run, making it general, accurate and highly efficient, with the ability to impact early stage PROTAC-based drug design campaigns where no structural information about the ternary complex structure is available.

## Introduction

In nature, Protein-Protein Interactions (PPI) have evolved to drive the assembly of macromolecular machines, impacting nearly all cellular processes^1,2,3^. For example, PPIs are critical to the functioning of systems such as ATP synthase^4^, ion channels^5^ or the ubiquitination machinery^6,7^. Among these, there is a clear distinction between obligate PPIs, i.e those in which the unbound monomers are unstable in vivo, and non-obligate PPIs^8,9^. Recently weak, transient, and ligand-mediated PPIs have gained attention. In particular, the field of PPIs mediated by Proteolysis Targeting Chimeras (PROTACS) has attracted significant attention and funding^10^. While the technology is more than 20 years old^11^, the base concept remains: a heterobifunctional PPI promoter composed of a ligand that binds to a protein target (warhead), a ligand that binds an E3 ligase (recruiter), and a flexible linker connecting the two ligands, is used to bring into close proximity a target and the ubiquitination enzyme^11–13^. In eukaryotes, the ubiquitin-proteasome system continuously degrades proteins^14^ to enforce quality control and to achieve the cell cycle. The PROTAC molecule hijacks the degradation pathway, promoting proximity-based protein ubiquitination. Therefore, the PROTAC facilitates the transfer of ubiquitin to a target protein, leading to poly-ubiquitination, and downstream degradation by the proteasome^13,15^. Some advantages of PROTAC-mediated degradation approaches include the use of lower compound concentration, decreased toxicity risks and the ability to target non-druggable proteins^16^.

In the last years, PROTAC-based approaches were pursued to some success, targeting and promoting the degradation of several proteins such as bromodomain 4 (BRD4)^16,17^, kinases^18,19^, fusion proteins^20,21^, growth factors^21,22^ and other proteins involved in various cancers^23,24^. Currently, at least 18 PROTACS are undergoing phase 1/2 clinical trials^25^. A compilation of bioactive PROTAC molecules, along with their description in terms of physicochemical properties was recently published and is expected to help campaigns of PROTAC drug design^26^.

While more and more experimentally determined structures for E3 Ligase-PROTAC-Target systems are being deposited in the Protein Data Bank (PDB)^27^, the data is still sparse for PROTAC design approaches. Thus, computational methodologies are being explored to predict the structure of ternary complexes and, using machine learning methods, degradation rates^28–32^. One of the most popular tools for PPI prediction is protein-protein docking, where the prediction depends on two critical steps: sampling of the conformational landscape of the protein-protein complex and scoring of the near-native poses obtained^33^. Most often, PPI docking protocols rely on rigid-body approximations, disregarding conformational flexibility or approximating it in a rough manner^34–40^. In the case of PROTAC-mediated ternary complexes, molecular docking approaches may require careful set-up and additional post-processing steps.

Indeed, recent approaches employ molecular docking combined with several steps of structure refinement and explicit inclusion of PROTAC ligand^30,31^. For example, the study by Weng and co-workers introduced a workflow (PROTAC-Model) which combines a local docking grid searching in rotational space algorithm (FRODOCK)^41^ with filtering steps, full PROTAC construction, ternary complex rescoring and a final optimization using RosettaDock^42–44^. One alternative route was proposed by Dixon and co-workers^45^, where a combination of molecular modeling, SAXS, and HDX-MS experiments led to the determination of physiologically relevant ternary complexes. One of the main points from this study is that the free energy landscape of these systems is not characterized by a single low-energy conformation but instead by several, interconvertible, low-energy minima, highlighting the complexity and flexibility of PROTAC-mediated biomolecular systems^45^. Finally, given the immense success that AlphaFold2 (AF2)^46^ and other machine learning-based methods have shown in the last CASP14 and CASP15 editions^47,48^, some researchers have pointed out that in the absence of experimental structures, future PROTAC-based drug design projects will use AF2 predicted structures^49^.

In this study, we present a general approach to generate ligase-target binding modes compatible with PROTAC-mediated ternary complexes determined by X-ray crystallography. Our aim is to produce a workflow that facilitates the initial stage of PROTAC *in silico* design, where the PROTAC molecule is still unknown or where structural information of the ternary complex is lacking. Our approach is tested on a manually curated dataset of 13 ternary complexes extracted from the PDB and relies on the predictions provided by LightDock^38,40,50^. LightDock is combined with energy rescoring afforded by VoroMQA^51^ and a filtering step based on the minimal solvent-accessible surface distance (SASD) between anchor atoms of the ligands bound to the ligase and target proteins, computed using Jwalk^52^. We show that restrained-LightDock simulations produce reasonable PPI structures, which are compatible with experimental structure of the ternary complex, for nearly all systems under scrutiny. We further compare the performance of our method with AlphaFold2-Multimer^46,53^, highlighting critical limitations in the machine-learning based method for this kind of application, and benchmark its computational efficiency against PROTAC-Model^31^ highlighting that the present approach is able to efficiently generate reasonable starting ligase-target binding modes.

## Methods

### Dataset selection and curation

Thirteen PROTAC ternary complex crystal structures were extracted from the PDB^27^, selected based on three criteria: (1) that the crystal structure contained an E3 ligase, a ligand molecule, and a target; (2) that the crystal structure resolution was below 4 Å and (3) that the ligand was a PROTAC and not a molecular glue. The dataset is composed of three types of E3 ligases (9 VHL^17,54–59,^ 2 Cereblon^60–63^ and 2 cIAP^64^), multiple different PROTAC molecules and a diverse set of protein targets, from kinases to bromodomains or proteins implicated in DNA repair (as WRD5). For the realistic docking case scenario, unbound monomers were selected to be different from the initial 13 crystal structures in terms of interface side-chain packing^54,64–71^. All protein termini were capped using the psfgen tool in VMD^72^, residue names were converted to standard names, and hydrogens were removed prior to molecular docking experiments. A listing of the structures used in the redocking and realistic experiments is given on **Tables S1** and **S2**.

### Molecular Docking and refinement protocol

Docking experiments were carried out using version 0.9.2 of the LightDock framework^38,40^. LightDock is a multiscale framework for the 3D determination of binary macromolecular complexes based on the Glowworm Swarm Optimization algorithm^73^. The sampling in LightDock is driven by the *swarm* approach, which relies on the metaphor that, in nature, *glowworms* (which represent ligand, or protein target, poses) feel attracted to each other depending on the amount of emitted light (scoring, energetic value of a docking pose). In this way, the docking poses, which constitute the *swarm* of *glowworms* in LightDock, are optimized towards the energetically more favorable ones through the translational, rotational, and, if the option is selected, some degree of flexibility at the level of the protein backbone is achieved through the optimization of an anisotropic network model (ANM)^38,40^. In principle, the ANM helps finding better docking poses since it “smoothes” the energetic docking search landscape. In this study, the ANM option was not explored for two reasons: (i) as the difference in RMSD between the individual monomers and the respective structures in the ternary complex crystal is small or non-existent (for the realistic docking or the redocking experiments, respectively), the ANM may actually just add noise and (ii), it would significantly increase the computational cost of each run because it would require that each docking simulation be run for at least another 50 steps to allow the optimization in ANM space to converge. Every swarm represents an independent docking simulation, and the number of swarms is automatically determined depending on the receptor surface nature, ensuring full sampling coverage of the protein complex surface. If contact residue-restraints are provided, swarms are filtered according to their compatibility with these restraints, focusing sampling to the region of interest described by the restraints^38,40^. To define the contact residues for restraints, the user specifies two lists of residues, one for the ligand (target protein) and one for the receptor (E3 ligase). Given a structure, a contact between two residues *i* and *j* is present if the distance between any heavy atom of *i* and any heavy atom of *j* (for the DFIRE scoring function) is below 3.9 Å. For each pose, we count the number of “satisfied residues’’ for the ligand list and the receptor list, where a satisfied residue in the ligand list is a residue which is in contact with any residue of the receptor list, and vice-versa. The number of satisfied residues for the ligand and the receptor are normalized by the total number of residues in the user lists, which leads to two fractions PL and PR, respectively. The fractions are combined with the docking score as follows:

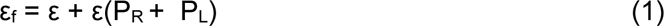

where ε is the unbiased score and εf is the final score. The term involving the satisfied fractions acts as a “booster”, promoting the poses which satisfy the most of the user-specified residues, as evaluated and benchmarked in different scenarios in ^40^. The number of glowworms per swarm was set to 200 (which is the default in the protocol), and 50 docking iterations were carried out (since 50 steps is generally enough for convergence if ANM is not enabled) using the DFIRE scoring function^40,74,75^.

The docking experiments were carried out in two scenarios: (1) redocking (or bound) and (2) realistic (unbound) docking, starting from the monomers within crystal ternary complexes and from independent monomeric forms of the proteins found in the PDB, respectively. Furthermore, in both redocking and realistic docking experiments, we explored the enforcing of contact residue restraints to help drive the docking towards the PPI. Contact residue restraints were defined by identifying residues in the ligand binding site whose C-alpha atoms were within a given distance (4, 6 or 8Å) from any heavy atom of the ligands bound either to the E3 ligase or the target protein, determined using MDAnalysis^76,77^. We excluded residues whose relative Solvent Accessible Surface Area (SASA) was below 25%, with SASA computed using naccess^78^. From each docking experiment, the top 200 poses were extracted according to the LightDock predicted ranking. Lightdock, like many other state-of-the-art docking approaches, does not sample side-chains due to the massive computational cost associated with sampling the configurational space comprising all accessible rotameric states of all amino acid side chains. While it would be possible to include backbone flexibility in the simulations via the ANM, for the reasons described above (computational cost of doubling the number of simulation steps to ensure convergence, possibility that the ANM would only add noise) we did not pursue this avenue. As such, docking experiments were carried out in a rigid-body fashion.

To improve protocol accuracy, a rescoring step was pursued using the energy function from VoroMQA^51^, which was sufficient in the case of the redocking experiment, and then filtered according to either the Center-of-Geometry (COG) distance between the recruiter and warhead ligands or the existence and length of the SASD between small-molecule anchor atoms^52^ for the realistic docking experiment. The former was computed using MDAnalysis^76,77^ while the latter was computed using Jwalk^52^. The SASD filter was calibrated using the predicted PPIs, exploring several SASD ranges (**Table S6**) and the reference structures in the benchmark dataset with the aim of recovering as many acceptable solutions as possible while also encompassing at least 90% of the SASDs computed on the crystal structures (see **Table S1**). The rescoring step corresponds to a re-ranking of the LightDock-generated structures in ascending order of the calculated VoroMQA energies. The filtering step shifts all solutions which do not pass the criteria to the bottom of the ranking, maintaining the order obtained from the energy rescoring. This filtering allows one to discard predicted PPI structures where the either or both of the binding sites are occluded, effectively acting as an implicit “PROTACability filter”.

### Docking accuracy

After each experiment, the accuracy of the docking protocol was evaluated using the CAPRI standards^79,80^ and the DockQ score^81^. These criteria employ three metrics between docking poses and crystallographic complexes, classifying docking solutions as incorrect, acceptable, medium or high quality. The three similarity measures are the fraction of conserved native contacts (Fnat), the interface root mean squared deviation (RMSD) and the ligand RMSD, where ligand refers to the target protein, which were computed in this study using C-alphas. As PROTAC-mediated ternary complexes are more and more suspected to be flexible^45^, we consider that any solution with acceptable or higher quality is a reasonable solution. The reference structures used for both experiments were the PROTAC ternary complex crystals. Docking accuracy was measured by evaluating the presence of solutions with at least acceptable quality in the top 1, 5, 10, 20, 50 or 100 protein-protein docking poses. Acceptable solutions thus adhere to either (i) a DockQ score ≥ 0.23, or (ii) a fraction of native contact (Fnat) >=10% and either ligand RMSD ≤ 10 Å or interface RMSD ≤ 4 Å^81^. A pose is acceptable if it passes either the CAPRI or DockQ criteria.

### AlphaFold-multimer structure predictions

Calculations using AlphaFold2-multimer^46^ were carried out using a local installation of version 2.1. All templates available on the PDB up to the 30^th^ of April 2018 were included. The AF2 confidence score was extracted to evaluate the confidence of the tool on the predicted protein-protein complexes^46^. The accuracy of AF2 predictions (**Figure S6**) was assessed using the same metrics as the predictions supplied by LightDock.

### PROTAC-Model benchmark calculations

To compare the results and performances of our method with PROTAC-Model^41^, we selected the systems shared between the two studies (**Table S5**) and ran PROTAC-Model using the default settings for a run without Rosetta refinement. The E3-ligase linker anchor atom selected for FRODOCKs’ local docking was the one present in the corresponding reference crystal structure. Calculations were carried out using 20 cores on a 40 CPU Intel Xeon Silver 4210R CPU machine running at 2.40 GHz.

## Results and discussion

### The docking workflow recovers PROTAC-mediated ternary complex-compatible PPIs

To evaluate the accuracy of our protocol, we first carried out redocking experiments on the benchmark dataset of 13 PROTAC-mediated complexes (**Table S1**). Here, we consider that the docking was successful if it generated at least one acceptable or higher quality solution within a given cohort (Top 1, 5, 10, 20, 50 or 100 ranked poses). From the information gathered in the literature, in particular the reports about ternary complex flexibility from Dixon and co-workers^45^, we hypothesized that accurate prediction of these interfaces would be a challenge for an unbiased docking algorithm. A way to help drive the docking towards the correct interface is through the introduction of contact residue restraints, which we pursued as detailed in the experimental section. This restraint selection approach ensures that the docking algorithm is directed towards the PPI while maintaining the overall protocol general and applicable to any ligase-target pair of interest. Importantly, these restraints correspond to an energetic bias which is added to the overall score per satisfied restraint. The redocking experiment is not realistic, as the side-chains are perfectly packed in the crystal, but represents the first test to assess protocol accuracy. **Figure 1** shows, for the 6BOY complex (**A**), a visual representation of the residues used in the restraints for the CRBN ligase at each selection threshold (**B**). **Table S3** contains an exhaustive description of the number of restraints per protein and system. As expected, unbiased docking simulations were not able to generate acceptable poses for the large majority of cases. For instance, we obtained an accuracy of 23% for the cohort of best 20 poses (**Figure 1C**, Top 20 light-blue bar). The reason behind the poor accuracy of the unbiased docking experiment is that the algorithm attempts to maximize the interaction surface as these are the most energetically favorable in the absence of restraints guiding the algorithm (**Figure S2**). On the other hand, the restrained docking calculations achieved satisfactory accuracy for the cohort of Top 20 poses (77%, 69% and 54% for calculations employing 4, 6 and 8 Å contact residue restraints, **Figure 1C**). Furthermore, we observed significant improvements in accuracy once the VoroMQA energy rescoring was carried out (**Figure 1D**). In particular, the protocol employing 6 Å contact residue restraints achieved 92% accuracy for the Top 20 cohort.

**Figure 1.**
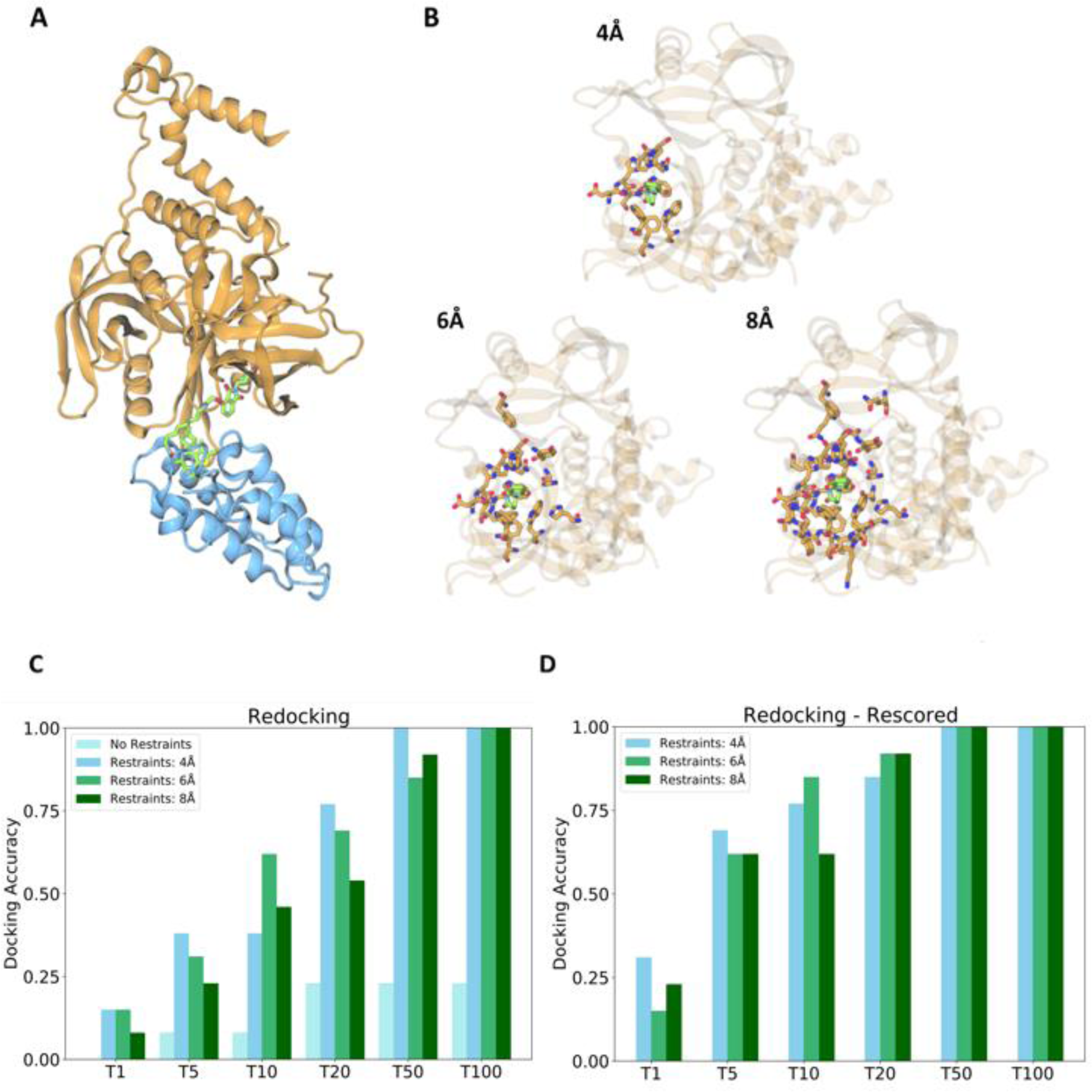
A) Crystal structure of the PROTAC-mediated complex 6BOY. The ligase component cereblon (CRBN, orange) is bound to the bromodomain target BRD4 (blue), with the complex being stabilized by the PROTAC ligand dBET6 (green). B) The center-of-mass of the thalidomide-like CRBN ligand from 6BOY (green) is used to select surface-accessible residues to act as docking restraints (orange) at different cutoff radii. C) Redocking calculations performed either with or without contact residue restraints. D) Redocking calculations after rescoring with the VoroMQA energy function.

### Minimal solvent-accessible path is essential to recover acceptable Ligase-Target binding poses in realistic docking scenarios

The same protein pairs that were used in the benchmark described above, were adopted for a realistic docking experiment. In this case, the docking inputs correspond to crystallographic ligand-bound structures of the monomeric forms for the E3 ligase and target proteins. Importantly, we did not enforce any prior knowledge on which linker composition and length might fit the ternary complex. As expected, there is a significant drop in accuracy when comparing the redocking with the realistic experiments (from 77% to 23% for the best performing setups at the Top 20 level, respectively), as illustrated by **Figures 2C** and **2D**. One possible reason may be that the sidechain packing in the redocking experiment is optimal and thus finding appropriate solutions is possible whereas in the realistic case, optimal side-chain orientations are not accessible. Nonetheless, we are able to prioritize more acceptable poses after VoroMQA energy rescoring^51^, leading to an increase in accuracy, from 23% to 46% at the Top 20 level for the best performing protocol (restraints defined with a 6 Å cutoff). To further improve the accuracy of the realistic docking calculations, we decided to impose a filtering scheme to remove highly ranked but physically unreasonable poses, where the COG euclidean distance between ligands (**Figure 2A**) is too large and/or the ligand binding sites are occluded. To account for binding site occlusion, we filtered the predicted docking poses through the existence and length of the solvent-accessible surface distance between the two ligands (SASD, **Figure 2B**).

**Figure 2.**
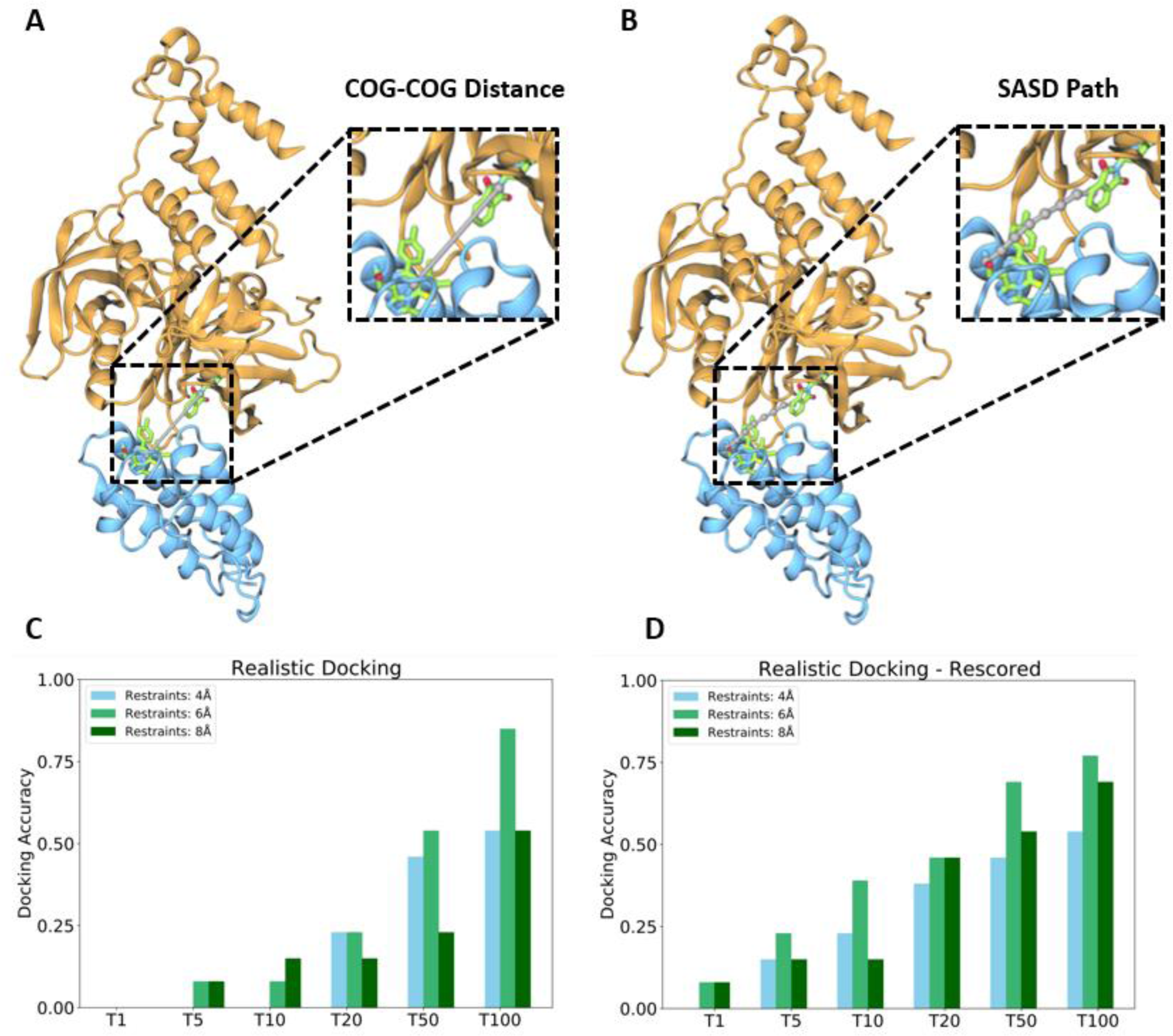
Realistic docking experiments carried out for the PROTAC benchmark dataset. A) Depiction of the COG-COG euclidean distance (grey) between the two ligands composing the dBet6 PROTAC (green), bound to the E3 ligase (orange) and a bromodomain target (blue) [PDB id: 6BOY]. B) Illustrative figure highlighting the minimal SASD path (grey) between the two ligands. C) Accuracy of realistic docking with contact residue restraints. D) Accuracy of realistic docking followed by VoroMQA rescoring. In the two bottom panels, the top scoring structures of each cohort (Top 1, 5, 10, 20, 50 or 100) were analyzed. Thus, T1 indicates that we analyzed the best scoring structure for all systems to see if its quality was at least acceptable according to either the CAPRI classification or the DockQ score.

The selected SASD filter was an interval from 3 to 13.7 Å (**Table S1**). As shown in **Figure 3A**, the filters improved remarkably the performance of the workflow, with the accuracy rising from 46% to 77% (Top 20, 6 Å contact residue restraints). However, the enhancement in performance observed here is still a product of a dual effect: the docking algorithms’ ability to produce meaningful solutions and the accuracy of the post-processing procedure, i.e. the energy-rescoring and SASD filtering. To ascertain the performance of the post-processing approach, we removed those systems for which no meaningful docking solutions were found for the best performing setup, which employs 6 Å contact residue restraints (**Figure 3B**), two VHL cases (PDB IDs 7JTP, 7KHH, **Figure S4**). In this analysis, we considered successes the cases where LightDock was able to generate at least one acceptable solution for that particular system, a classification which allows to decouple the docking engines’ ability to produce meaningful solutions and the post-processing filters’ ability to recover those solutions. Since the structural differences of the ligase-target systems between their ternary complex and monomeric states are small (see **Table S2**), and their interfaces in the ternary complex are shallow (and may be as small as two contact residues), we believe that even small perturbations to sidechain orientation may be sufficient to induce the algorithm into error. We found that after removal of those systems, the protocol exhibits an accuracy of 82% and 91% for the Top 10 and 20 poses, respectively. Interestingly, if we only consider the most well studied ligases (CRBN and VHL), discarding the two systems employing inhibitor of apoptosis E3 ligases (cIAPs), the accuracy of the procedure rises to 89% and 100% at the Top 10 and Top 20 levels respectively. In conclusion, we achieved good accuracy if one considers that unbound monomer docking in the case of PROTACs is challenging and no structural refinement was performed. As examples, the highest ranked structures found after PROTACability filtering for three complexes are given in **Figure 3C-E** whereas **Figure S3** illustrates the highest-ranking acceptable solutions for all systems here explored.

**Figure 3.**
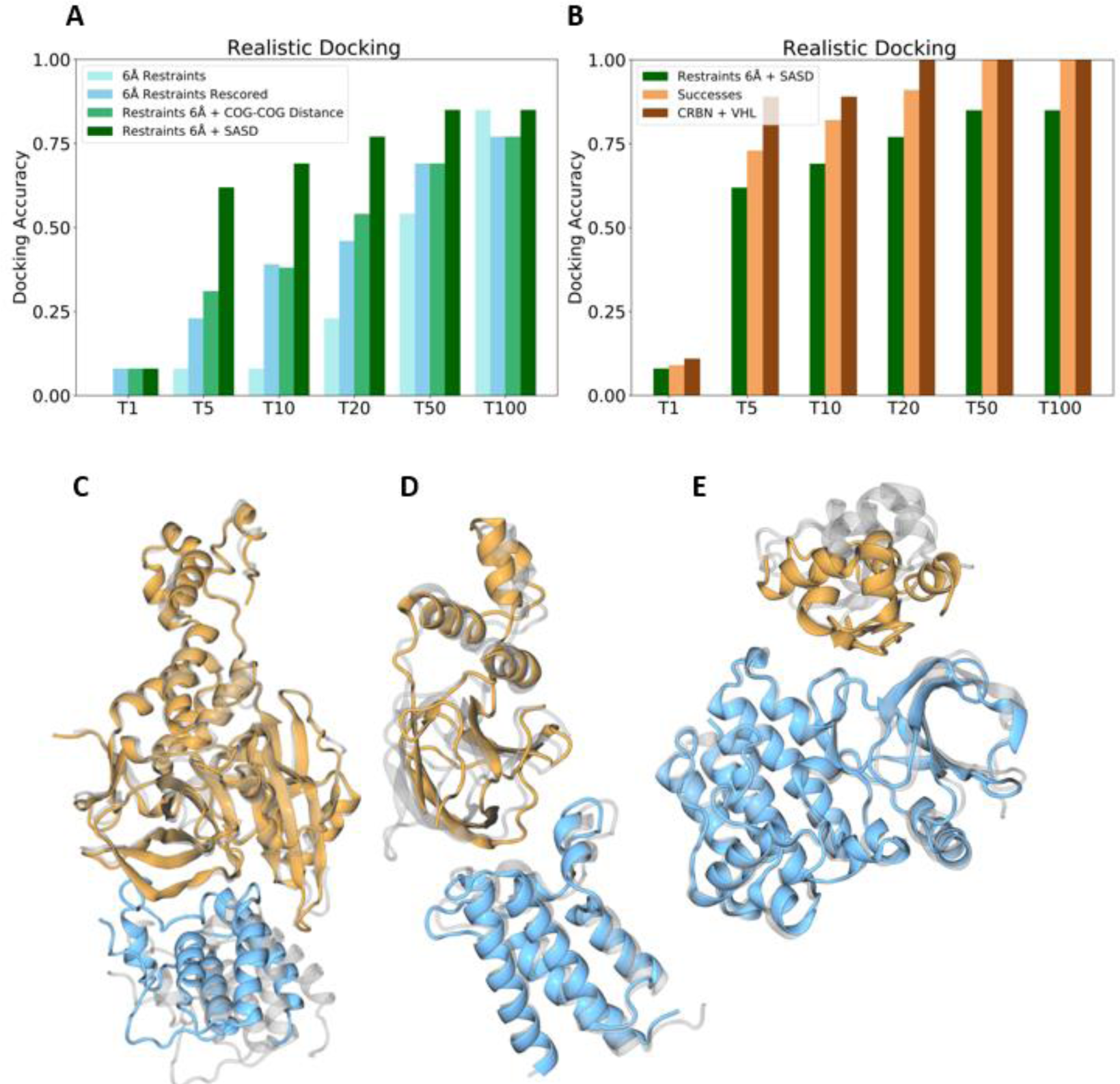
Accuracy of the realistic docking followed by rescoring and SASD filtering. A) Docking accuracy before and after filtering the predicted poses using either the COG-COG euclidean distance between ligands or the SASD computed using Jwalk. B) Summary of the performance of the best protocol. The first bar corresponds to the best performing setup, employing both VoroMQA energy rescoring and SASD filtering (dark green). The success cases are those (11 out of 13) for which at least acceptable-scored solutions were found by the docking engine in the full top 200 predicted structures (light orange). The last bar (dark brown) highlights the performance of the protocol when considering the successful cases that contain either CRBN or VHL ligases (9 out of 11 in total). C) Showcase comparing the 6BOY crystal structure (grey) to the best prediction (ranked first) obtained using our workflow (colored). D) Showcase comparing the 6HAX to crystal structure (grey) to the best prediction (ranked third) obtained using our workflow (colored). E) Showcase comparing the 6W8I crystal structure (grey) to the best prediction (ranked sixth) obtained using our workflow (colored).

## Discussion

In this study, we propose a novel workflow for the generation of PROTAC-compatible PPI structures in the case that no known PROTAC molecule exists, no information of a possible linker is available, and the ligase-recruiter and target-warhead co-crystals are available. To this end, we manually curated a dataset of 13 PROTAC-containing ternary complexes from the PDB to serve as a benchmark dataset. Using this dataset for protocol benchmarking, we carried out both redocking and realistic docking experiments (see **Results**) where the accuracy of the docking was measured by evaluating the quality of the predicted structures using CAPRI and DockQ criteria^79–81^.

We found that both the inclusion of contact residue restraints and the rescoring using the energy function of VoroMQA^51^, were fundamental to increase the accuracy of the calculations in both redocking and realistic docking experiments. In the case of redocking, the best performing protocol (restraints defined with a 6 Å cutoff, rescored) achieved a striking accuracy of 85% and 92% for the 10 or 20 best ranked predicted PPIs, respectively. The unbiased docking experiments, in comparison, yielded poor accuracy. This was expected, as most docking scoring functions are developed to reproduce interfaces with large contact areas and PROTAC-mediated complexes exhibit typically small and transient PPIs, as illustrated in **Figure S1**. In the realistic docking experiments, the accuracy was much lower, even with restraints (below 50% for the Top 20). Since many high-ranking solutions presented occlusion of the ligand binding sites in either one or both of the proteins, we envisioned that an extra filtering step was needed, such that the predicted PPIs where it is impossible to thread a solvent-accessible path connecting the two ligands would be discarded. Based on this idea, we decided to use the minimal SASD^52^ as a filter and applied it to the VoroMQA-rescored predicted PPIs. The application of the filter improves the docking accuracy, which rises from 46% to 77% for the Top 20 cohort. Nonetheless, the approach was unable to produce acceptable solutions in two cases. If we consider only the remaining 11 cases, we state that the post-processing workflow (rescoring + SASD filtering) is highly accurate for both the Top 10 and Top 20 cohorts (82% and 91% accuracy, respectively). Nonetheless, the recovery of acceptable solutions for the two cIAP systems comes at the cost of overall performance at the early stages (Top 5 to Top 20, **Figure 3E**). It is important to note that, recently, researchers have pointed out to the fact that the cIAP structures used in this work contain only one domain of the full cIAP protein and that the full length cIAP interacts with their substrates from different domains, which bind different parts of the substrate^84^, thus making a proper prediction of these PPIs challenging.

Our protocol is general and applicable to any ligase-target protein pair, as it only requires the corresponding ligand-bound structures of the monomeric forms for each protein. Furthermore, it is computationally cheap as only three steps are necessary: (1) restrained docking using LightDock, (2) energy rescoring using VoroMQA and (3) filtering by SASD. As such, our protocol may be useful to accelerate PROTAC-based drug discovery campaigns where no known PROTAC is available while being computationally tractable, with an average run-time per system of 24.1 minutes using 20 cpu cores (481 minutes per system on a single core (**Table S4**), and simple to employ.

Nonetheless, there are some inherent limitations to our work. Having taken the decision to keep it computationally cheap and fast, there are no structural refinement steps. This is clearly fundamental when looking at the showcases in **Figures 3C, 3D** and **3E**. It appears that even when guided by restraints, the LightDock still tries to maximize the interface contact area, leading to a super-packed binding mode. One of the reasons for this behavior is that the training set of the DFIRE^74^ scoring function was based on typical protein-protein complexes, which tend to have larger interface areas than those present in PROTAC-mediated ternary complexes (**Figure S2**). Possible avenues for improvement could pass by including protein flexibility by means of Normal Mode Analysis (NMA) in internal coordinates^82^, the inclusion of the ligand molecules in the structures during docking, testing other docking scoring functions or refinement of the docked poses based on Molecular Dynamics simulations. Finally, Jwalk^52^ does not consider the ligand molecules when computing the minimal SASD, meaning that this path may run across the ligand molecular structures depending on the positions of the anchor atoms. A further potential improvement can come from the application of an *in-house* developed tool which does exclude from the path construction the voxels occupied by ligand atoms (**Figure S5**). As such, the proposed workflow is accurate but improvable, and future work in our group is aimed at increasing the quality of the docking-predicted PPIs.

Other computational studies based on molecular docking have been pursued in recent years, aiming at the establishment of easy-to-use and accurate PROTAC drug design strategies. For example, the group behind PROsettaC^30^ and the group of Tingjun Hou^31^ both aimed at reproducing the crystallographic PPI of PROTAC-mediated ternary complexes. The workflow by Weng and co-workers^31^ was shown to be able to generate medium to high quality solutions for 12 out of the 14 PROTAC systems queried in a realistic docking scenario. However, it requires *a priori* knowledge of the PROTAC to use, which is critical for the filtering and retrieval of the protein-protein docking poses closest to the reference structure. We hypothesize that this is not the most optimal approach, since according to the data from Dixon and co-workers^45^, the crystal structure is one possible solution but it is not necessarily the only acceptable solution and trying to reproduce it may lead to discarding other biologically relevant conformations.

Nonetheless, we compared the computational cost and the accuracy of our method to PROTAC-Model by Weng et al^41^ (**Table S5**) across the systems shared between the two studies without employing PROTAC-Models’ Rosetta refinement, using 20 cores. From this comparison, we observe that while PROTAC-Model is superior to our approach at reproducing VHL-based ternary complex crystallographic structures, our method was (1) able to recover solutions for both cIAP systems and (2) performs slightly better than PROTAC-Model for the Cereblon systems studied. Furthermore, while our method took, on average, 18.7 minutes to generate the PPI predictions, PROTAC-Model takes, on average, over two hours (155.4 minutes), meaning that our approach is 8x more efficient that PROTAC-Model while using less information since our method does not require knowledge of the linker portion binding the two ends of the PROTAC.

Since DeepMinds’ AlphaFold2 is now able to predict protein-protein complexes in the multimer variant, we also questioned whether AF2-multimer^46^ would be able to correctly predict the PROTAC-mediated PPI interfaces. As shown in **Figure S7**, the AFs’ accuracy according to DockQ and/or CAPRI was low. As an example, the predictions obtained for 6BOY when run through AF2-Multimer, which are in complete disagreement with the experimental structure. A possible avenue to improve on these predictions is the inclusion of ligands in the modeling process, as proposed by Hekkelman and co-workers^83^. Thus, it appears that AF2- multimer cannot predict PROTAC-mediated PPIs, highlighting the critical contribution of the restraints. For the moment, physical-based approaches combined, as described here, seem to be a better choice towards computational design of PROTACs-compatible PPIs. Nonetheless, future work in our group will aim at improving the accuracy and capabilities of the PROTACability workflow by exploring different scoring functions (including free energy methods like the Molecular Mechanics-Poisson Boltzmann (or its Generalized-Born variant) Surface Area method^85^), protein flexibility and different filtering schemes.

## Conclusion

In this study we have described a new LightDock-based protocol for PROTAC drug design. Our workflow combines restraint-enabled LightDock simulations with energy-based rescoring from VoroMQA and SASD calculations from Jwalk to yield at least acceptable quality PPI complex structures, starting only from the ligand-bound monomeric form of the E3 ligase and the protein target. The protocol is generalizable to any E3 ligase/target pair, fast and accurate, achieving 92% and 77% accuracy for the top 20 cohort on the redocking and realistic docking experiments, respectively, when benchmarked against a manually curated and diverse dataset of PROTAC-mediated ternary complexes. In the case that no PROTAC molecule is known to bind the ligase/target pair of interest, our approach is able to provide initial configurations which reproduce the protein-protein binding interface and provide the minimal solvent accessible path connecting the ligand and target ligands, potentially accelerating PROTAC drug design campaigns by narrowing down the chemical space to be searched for linker selection. Compared to a state-of-the-art method, PROTAC-Model, our method is a compromise between accuracy and computational efficiency. While it is less accurate than PROTAC-Model at reproducing VHL-based ternary complex crystal PPIs, it is still able to generate near-native solutions, using a fraction of the computational time.

## Acknowledgements

Computer hours awarded to the GENCI project number 2022-A0120713456 on the Jean-Zay and Joliot-Curie Rome clusters of the French National Supercomputing Center are gratefully acknowledged. We gratefully acknowledge CC-IN2P3 (https://cc.in2p3.fr) for providing a significant amount of the computing resources and services needed for this work. The authors also acknowledge the support provided by the CNRS and PharmCADD.

## Supporting Information

Additional details, including a computational efficiency benchmark for PROTACability. the datasets used for model benchmarking, data on the parametrization of the SASD filter, comparison of Protein-Protein Interface sizes, and a schematic workflow of the PROTACability protocol are given.

## Data availability and code

A step-by-step tutorial, codes, scripts and an example using 6HAX (for realistic docking) are available on GitHub (https://github.com/GilbertoPPereira/PROTACability). All reference structures used in the dataset and results obtained in this work are available at Zenodo: https://zenodo.org/record/8136088.

---

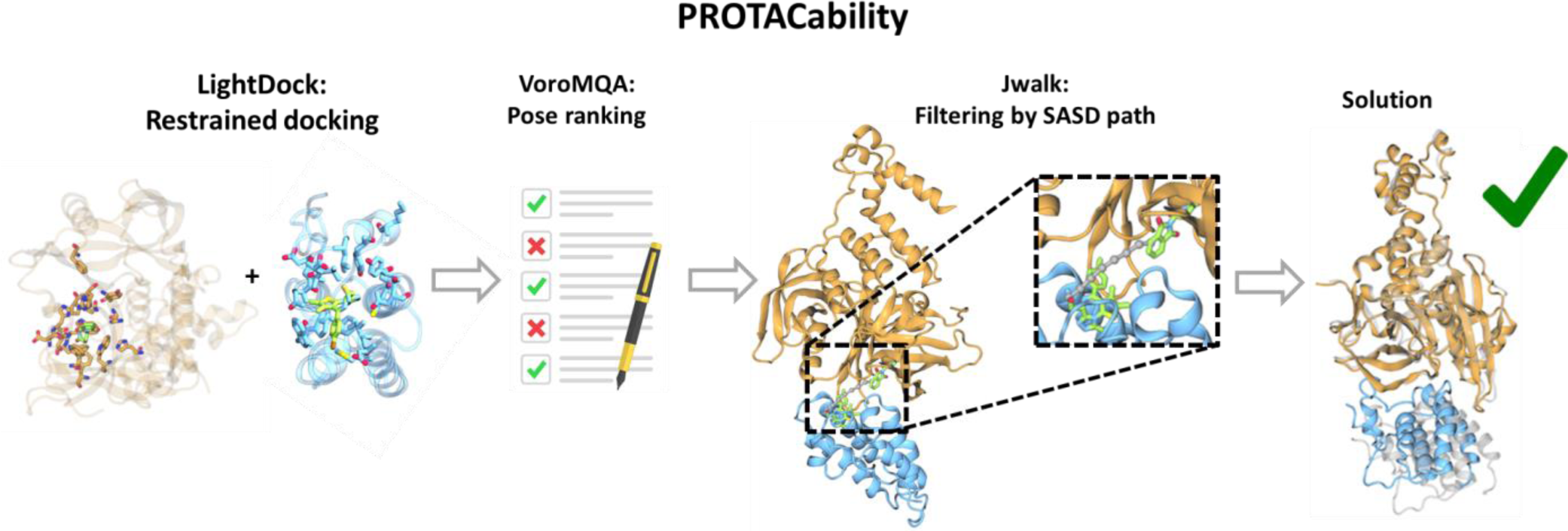

