## Supplemental Material for "Rational Prediction of PROTAC-compatible Protein-Protein Interfaces by Molecular Docking"

**Table S1:** Benchmark dataset used for protocol evaluation.

| Complex | Ligase | Target | PROTAC | Resolution (Å) | SASD (Å) |
|---|---|---|---|---|---|
| 5T35 | VHL | BRD4 | MZ1 | 2.7 | 9.9 |
| 6BN7 | CRBN | BRD4 | dBET23 | 3.5 | 11.9 |
| 6BOY | CRBN | BRD4 | dBET6 | 3.3 | 15.2 |
| 6HAX | VHL | SMARCA2 | PROTAC 2 | 2.4 | 7.1 |
| 6HR2 | VHL | SMARCA4 | PROTAC 2 | 1.8 | 7.4 |
| 6W7O | cIAP | BTK | compound 17 | 2.2 | 8.9 |
| 6W8I | cIAP | BTK | compound 15 | 3.8 | 13.5 |
| 6ZHC | VHL | Bcl-Xl | PROTAC6 | 1.9 | 8.1 |
| 7JTO | VHL | WDR5 | MS33 | 1.7 | 12.4 |
| 7JTP | VHL | WDR5 | MS67 | 2.1 | 6.5 |
| 7KHH | VHL | BRD4-BD1 | Compound9 | 2.3 | 6.6 |
| 7Q2J | VHL | WDR5 | Homer | 2.5 | 6.6 |
| 7PI4 | VHL | FAK | GSK215 | 2.24 | 8.3 |

CRBN: Cereblon; VHL: Von-Hippel Landau; SASD: Solvent Accessible Surface Distance computed using Jwalk[1].

**Table S2:** Monomers of the ligase/target pairs used in the realistic docking experiment.

| Complex | Ligase Monomer | RMSD (Å) | | | Target Monomer | RMSD (Å) | | |
|---|---|---|---|---|---|---|---|---|
| | | Full | Binding site | Binding site (Cα) | | Full | Binding site | Binding site (Cα) |
| 5T35 | 3ZUN | 1.2 | 1.7 | 0.8 | 7C6P | 0.9 | 1.2 | 0.5 |
| 6BN7 | 5V3O | 1.5 | 1.8 | 1.0 | 7C2Z | 1.4 | 1.3 | 0.7 |
| 6BOY | 5V3O | 1.6 | 1.7 | 1.1 | 7C2Z | 1.5 | 1.5 | 0.7 |
| 6HAX | 3ZUN | 1.2 | 1.8 | 0.7 | 6HAZ | 1.2 | 1.5 | 0.6 |
| 6HR2 | 3ZUN | 1.1 | 1.7 | 0.7 | 6ZS2 | 1.2 | 1.5 | 1.9 |
| 6W7O | 6W74 | 0.9 | 0.6 | 0.4 | 6W8I | 1.3 | 1.1 | 0.6 |
| 6W8I | 6W74 | 1.6 | 2.9 | 1.5 | 6W7O | 1.6 | 1.6 | 1.0 |
| 6ZHC | 3ZUN | 1.0 | 0.8 | 0.4 | 2BZW | 3.1 | 4.5 | 4.1 |
| 7JTO | 3ZUN | 1.1 | 1.6 | 0.4 | 2H9M | 0.9 | 1.1 | 0.6 |
| 7JTP | 3ZUN | 1.1 | 1.7 | 0.5 | 2H9M | 0.9 | 1.0 | 0.5 |
| 7KHH | 3ZUN | 1.1 | 1.2 | 0.4 | 7C2Z | 1.2 | 1.0 | 0.6 |
| 7Q2J | 3ZUN | 1.1 | 1.1 | 0.4 | 2H9M | 1.0 | 1.4 | 0.5 |
| 7PI4 | 3ZUN | 1.0 | 1.1 | 0.5 | 2ETM | 2.2 | 1.5 | 1.0 |

The RMSD measurements here described were carried out using ProFit[3], comparing the monomeric structures of the proteins to the corresponding protein in the crystal structure. The RMSD for the full monomer was measured using all heavy atoms. The binding site RMSD was measured either using only the alpha carbons or including the heavy atoms of the side-chains, on the set of residues that are at less than 10 Å from the contacting chain.

**Table S3:** Number of restraints per system and restraint cut-off.

| Complex | Redocking Rest: 4A | Redocking Rest: 6A | Redocking Rest: 8A | Realistic Rest: 4A | Realistic Rest: 6A | Realistic Rest: 8A |
|---|---|---|---|---|---|---|
| 5T35 | 27 | 36 | 61 | 23 | 37 | 62 |
| 6BN7 | 23 | 36 | 58 | 20 | 33 | 58 |
| 6BOY | 24 | 34 | 58 | 18 | 31 | 58 |
| 6HAX | 30 | 43 | 59 | 27 | 39 | 60 |
| 6HR2 | 31 | 45 | 66 | 29 | 40 | 64 |
| 6W7O | 32 | 47 | 75 | 28 | 46 | 71 |
| 6W8I | 33 | 47 | 71 | 28 | 44 | 69 |
| 6ZHC | 32 | 45 | 71 | 26 | 38 | 65 |
| 7JTO | 33 | 53 | 73 | 33 | 47 | 72 |
| 7JTP | 33 | 49 | 75 | 33 | 48 | 74 |
| 7KHH | 24 | 33 | 57 | 30 | 41 | 65 |
| 7Q2J | 34 | 53 | 75 | 30 | 46 | 71 |
| 7PI4 | 34 | 47 | 73 | 27 | 49 | 77 |

**Table S4:** Performance of the workflow in minutes running on a single CPU core**.**

| System | LightDock | | | Post-Processing | | | Total |
|---|---|---|---|---|---|---|---|
| | Docking | Pose Generation | Pose Clustering | DockQ | VoroMQA | Jwalk | |
| 5t35 | 260 | 5 | 6 | 2 | 15 | 3 | 291 |
| 6bn7 | 300 | 5 | 7 | 3 | 27 | 4 | 346 |
| 6boy | 360 | 5 | 8 | 3 | 28 | 4 | 408 |
| 6hax | 280 | 6 | 10 | 2 | 16 | 3 | 317 |
| 6hr2 | 320 | 7 | 9 | 2 | 16 | 3 | 357 |
| 6w7o | 380 | 6 | 7 | 2 | 20 | 3 | 418 |
| 6w8i | 400 | 5 | 7 | 2 | 20 | 3 | 437 |
| 6zhc | 360 | 6 | 9 | 2 | 17 | 3 | 397 |
| 7jto | 660 | 9 | 12 | 3 | 25 | 4 | 713 |
| 7jtp | 660 | 9 | 11 | 3 | 25 | 4 | 712 |
| 7khh | 360 | 6 | 10 | 2 | 18 | 3 | 399 |
| 7pi4 | 600 | 9 | 12 | 3 | 23 | 3 | 650 |
| 7q2j | 760 | 10 | 12 | 3 | 25 | 4 | 814 |
| Average | 438 | 7 | 9 | 2 | 21 | 3 | 481 |
| SD | 162 | 2 | 2 | 1 | 4 | 0 | 168 |

All run-time data is provided for a single core; S.E.M is the standard error of the mean associated with the average run-time of the workflow in minutes.

**Table S5:** Comparison to the PROTAC-Model code.

| | PROTAC-Model | | PROTACability | |
|---|---|---|---|---|
| **SYSTEM** | **Run-time (min)** | **Quality** | **Run-time (min)** | **Quality** |
| 5T35 | 161 | High | 15 | Acceptable |
| 6BOY | 135 | Acceptable | 17 | Medium |
| 6BN7 | 155 | Acceptable | 20 | Acceptable |
| 6HAX | 173 | Medium | 16 | Acceptable |
| 6ZHC | 73 | Medium | 18 | Acceptable |
| 6HR2 | 311 | Medium | 20 | Acceptable |
| 6W7O | 17 | Medium | 21 | Acceptable |
| 6W8I | 268 | No Solution | 22 | Acceptable |
| 7KHH | 106 | Acceptable | 20 | Acceptable |

All calculations were carried out using 20 CPU cores, on an Intel Xeon Silver 4210R CPU machine running at 2.40 GHz.

**Table S6:** Selection of SASD Filter range

The SASD filter is a general filtering scheme which is defined as the solvent-accessible distance path between the achor atoms of the recruiter and warhead ligands within a predicted Ligase-Target pose. It was constructed to remove unphysical predictions produced by LightDock (such as those poses where one or both ligand binding sides were occluded and/or a path between the two ligands could not be constructed). It was applied to all complexes, and was calibrated such that it covered at least 90% of the SASD values from the crystal references while maximizing the accuracy of the protocol for the realistic docking experiment. Indeed, the only system whose SASD value falls outside the filter is 6BOY. The accuracy of the protocol with varying selection thresholds is shown below:

| **Cohort** | **3[a]-12[b]** | **3-12.5[b]** | **3-13[b]** | **3-13.5[b]** | **3-13.7[b]** | **3-14[b]** | **3-14.5[b]** | **3-15[b]** | **3-15.3[b]** | **3-15.5[b]** |
|---|---|---|---|---|---|---|---|---|---|---|
| Top1 | 0.08 | 0.08 | 0.08 | 0.08 | 0.08 | 0.08 | 0.08 | 0.08 | 0.08 | 0.08 |
| Top5 | 0.62 | 0.62 | 0.62 | 0.62 | 0.62 | 0.62 | 0.62 | 0.54 | 0.54 | 0.54 |
| Top10 | 0.62 | 0.62 | 0.62 | 0.62 | 0.69 | 0.69 | 0.69 | 0.69 | 0.69 | 0.69 |
| Top20 | 0.69 | 0.69 | 0.69 | 0.69 | 0.77 | 0.77 | 0.77 | 0.77 | 0.77 | 0.77 |
| Top50 | 0.77 | 0.77 | 0.77 | 0.77 | 0.85 | 0.85 | 0.85 | 0.85 | 0.85 | 0.85 |
| Top100 | 0.77 | 0.77 | 0.77 | 0.77 | 0.85 | 0.85 | 0.85 | 0.85 | 0.85 | 0.85 |

a - Lower limit of the SASD range in Å; b - Higher limit of the SASD range in Å; Top1 - Evaluation of accuracy (as a fraction of number of successful systems/total number of systems studied) considering the best scored predicted ligase-target structures with respect to a crystallographic reference. Top5 - Evaluation of accuracy (as a fraction of number of successful systems/total number of systems studied) considering the five best scored predicted ligase-target structures with respect to a crystallographic reference.

As is observable from the Table, the range which maximizes the accuracy of the protocol in the early rankings is between 3-13.7 and 3-14.5 Å. More than that, and we see a drop in accuracy at the Top5 cohort. We consider the protocol to be linker independent because the filtering scheme is based on whether it is possible to connect the anchor atom of each ligand through a solute-free volume such that a PROTAC can be constructed and the only assumption it makes is that the linker must not be too long. Indeed, it has been shown previously that too long linkers in PROTAC molecules can lead to suboptimal degradation[10]. No assumption on the bulkiness, chemical composition or flexibility is made, such that within a range of SASD, any linker could be constructed. The filtering scheme here is, nonetheless, dependent on the “training set” used and thus, as more ternary structures are published, it may be re-run and adjusted to be more general.

**Figure S1:** Comparison of Protein-Protein Interface sizes between PROTAC-mediated ternary crystal structures, AlphaFold2-Multimer models, and solutions obtained from the unbiased redocking experiments using LightDock.

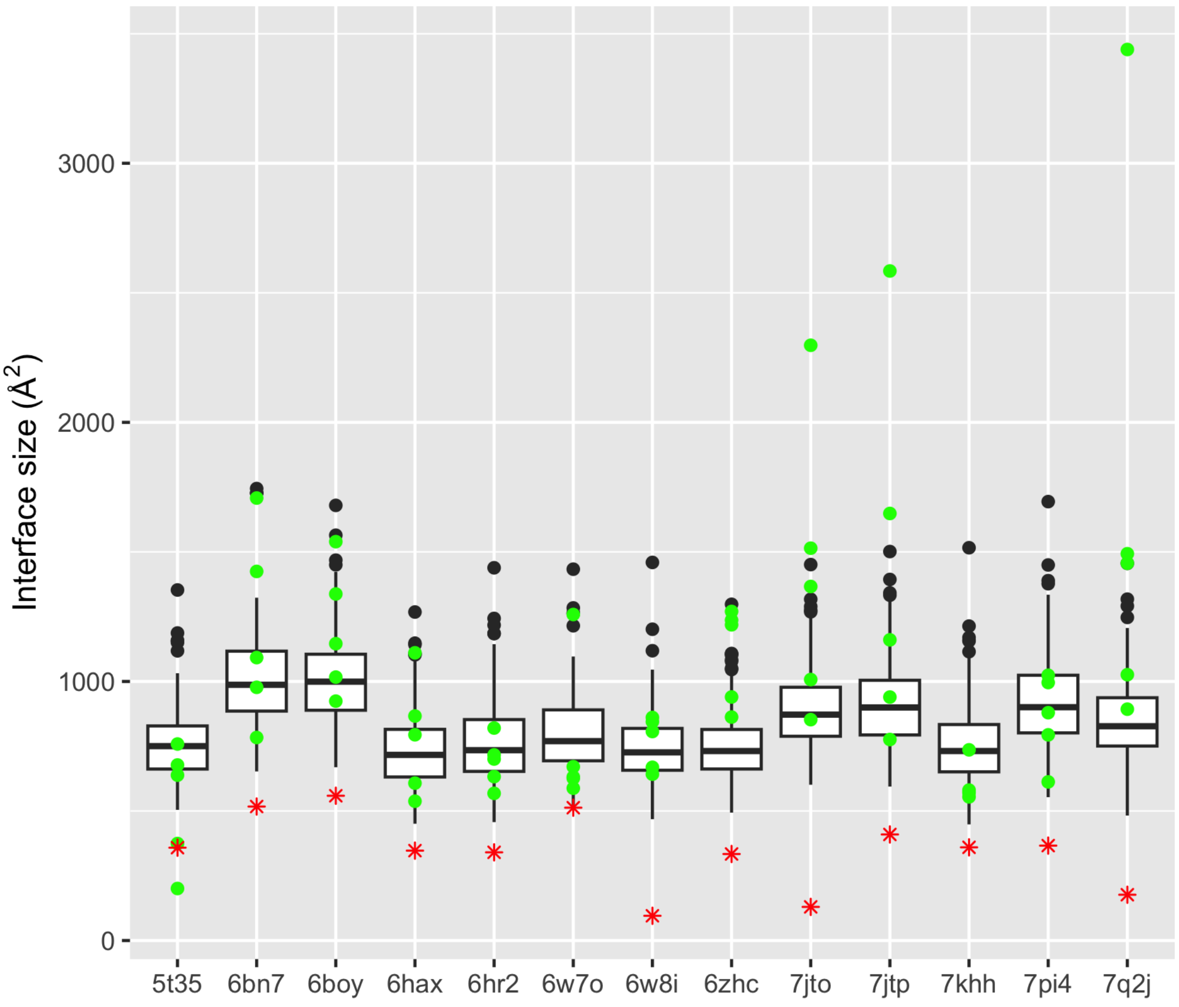


**Figure S1** - Comparison between interface sizes of the PROTAC-mediated ternary crystal structures (red), AlphaFold2-Multimer models[2] (green), and solutions obtained from the unbiased redocking experiments using LightDock (black).

The overlap between the protein-protein interface of solutions from LightDock[4–6] and models from AlphaFold2-Multimer highlights that both the training set of the DFIRE[7] scoring function and that of the much more recent AI-based model are heavily biased towards protein-protein complexes who have large interfaces and thus may be unsuited, on their own, to tackle small, ligand-mediated and transient PPIs.

**Figure S2:** Comparison between Protein-Protein interface sizes of PROTAC-mediated ternary complexes and interfaces in the Kundrotas dataset of protein dimers.

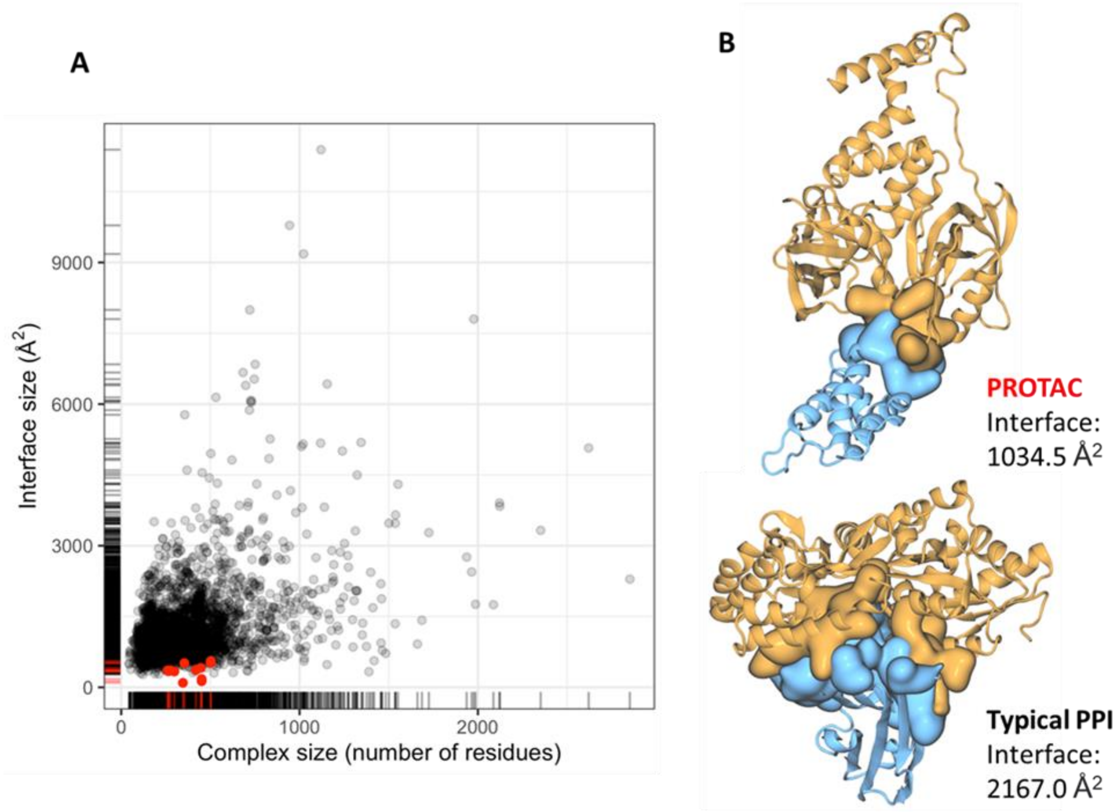


**Figure S2** - A) Distribution of protein-protein interfaces from the ternary and the reference data set. The size of the protein-protein interface (y axis) is plotted against the size of the protein complex (x axis) for a non-redundant set of 2552 hetero-dimers (background; in black) and for the 13 target/ligase/ligand complexes after removing the ligand (in red). Interface size is measured by the average change of surface accessibility per protein partner; complex size is measured by the sum of the number of residues in the two protein partners. B) Visual example of the interface area from a typical PPI and a PROTAC-mediated complex with the same number of aminoacid residues (6BOY and 5M14 PDB IDs, respectively).

We compared the size of ternary complex interfaces (13) with respect to a standard dataset composed of 2552 protein dimers[8]. The interface size was computed for a set of 2552 non-redundant heterodimers and for the dataset of PROTAC ternary complexes using the naccess tool[9]. The interface size change upon binding was estimated by subtracting the surface accessible area of each protein partner to that of the complex and divide the difference by two. Comparison between the two datasets was performed by evaluating complex size in terms of number of amino acids and interface area, excluding the PROTAC ligand from the ternary complexes beforehand. As shown in **Figure S2A**, the average interface area of the ternary dataset complexes is much smaller than that of the reference set, with interface sizes between 96 and 654 Å$^2$, for a total complex size between 260 and 675 residues. This result was expected, since these PPIs are, in a sense, transient and appear to be stabilized upon PROTAC binding. This suggests that PROTAC-mediated ternary complex interfaces are transient and rely on the PROTAC to form instead of relying on maximizing the interaction network between the protein partners, as illustrated in **Figure S2B**.

**Figure S3:** Top ranked predicted structures obtained in the realistic docking experiments.

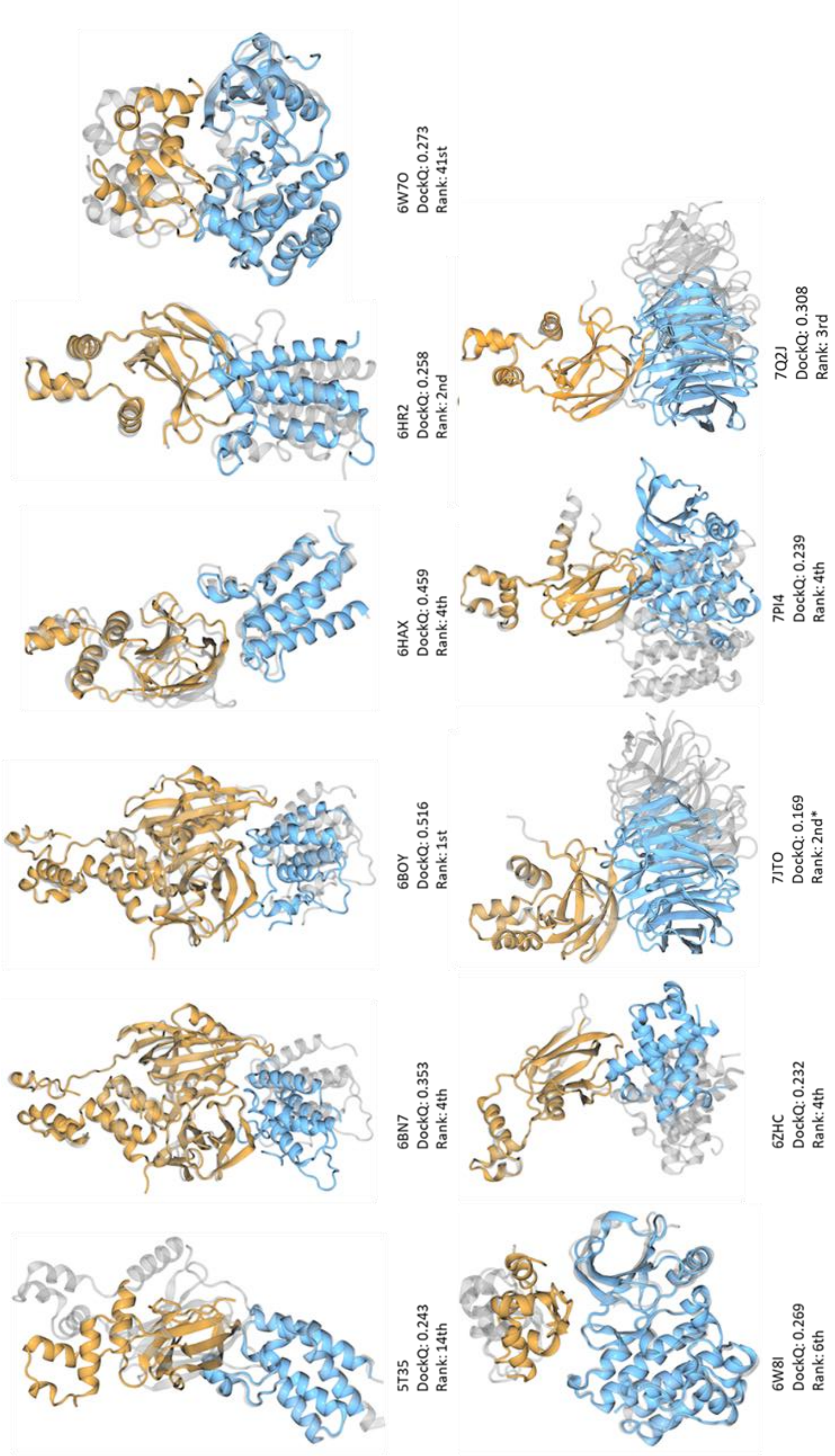


**Figure S3 -** Best solutions obtained in the realistic docking experiments. The reference structure (Grey) is overlaid with the best predicted PPI (colored, orange and blue represent the ligase and target proteins respectively), after filtering each of the 200 energy-rescored solutions with the minimal SASD cut-off. Although the 7JTO system fails the DockQ criteria (DockQ < 0.23), it passes the CAPRI rules with a Fraction of Native contacts (Fnat) = 0.25 (Minimum Fnat > 0.1) and an interface-RMSD (i-RMSD) of 3.484Å (Minimum i-RMSD < 4Å).

**Figure S4:** Top ranked predicted structures obtained in the realistic docking experiment for systems where no acceptable or higher quality solution was kept

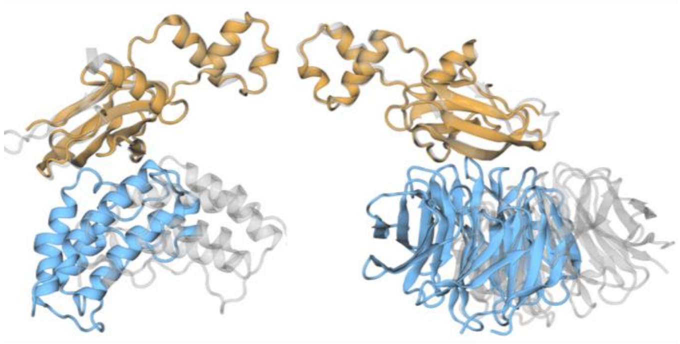

7KHH
DockQ: 0.117
Rank: 1st

7JTP
DockQ: 0.060
Rank: 1st

**Figure S4 -** Best solutions obtained in the realistic docking experiment for each system where no docking solution was found. The reference structure (Grey) is overlaid with the best predicted PPI (colored, orange and blue represent the ligase and target proteins respectively), after filtering each of the 200 energy-rescored solutions with the minimal SASD cut-off.

**Figure S5:** Comparison between the paths computed using Jwalk and Linky, a under-development tool for building optimal SASD paths.

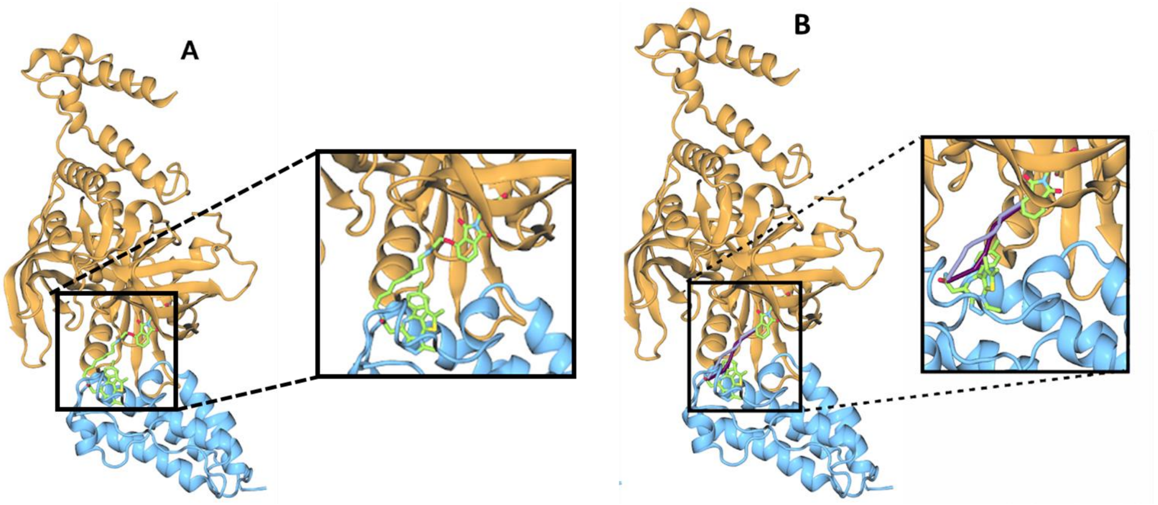


**Figure S5 -** Different methods to compute the minimal SASD path. A) Reference structure of 6BOY (ligase in orange; target in blue) with PROTAC (green) bound. The inset highlights the PROTAC-mediated PPI; B) Comparison between the SASD paths built by Jwalk[1] (purple) and linky (developed in-house, in mauve). The inset shows that the Jwalk path overlaps with ligand atoms (as Jwalk is ignorant to the volume occupied by the ligands) and that linky builds a more realistic path by also considering ligands solvent excluded volume.

**Figure S6:** PROTACability workflow.

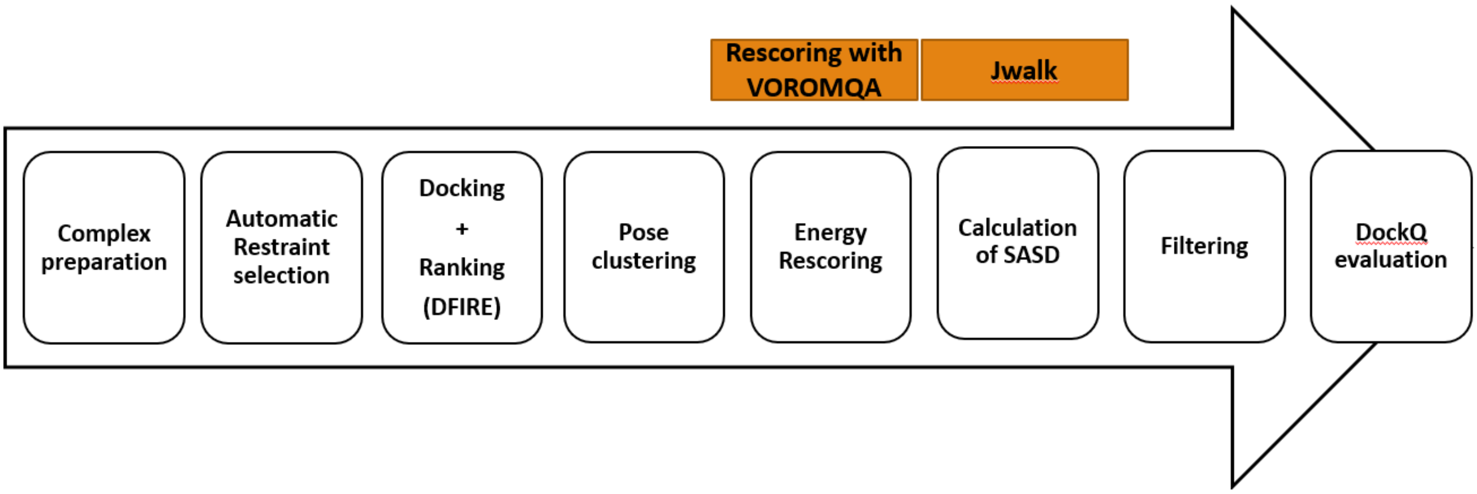


**Figure S6 –** Proposed workflow for PROTACability estimation of predicted PROTAC-mediated Protein-Protein Interfaces.

**Figure S7:** AlphaFold2 is unable to predict PROTAC-mediated Protein-Protein complexes

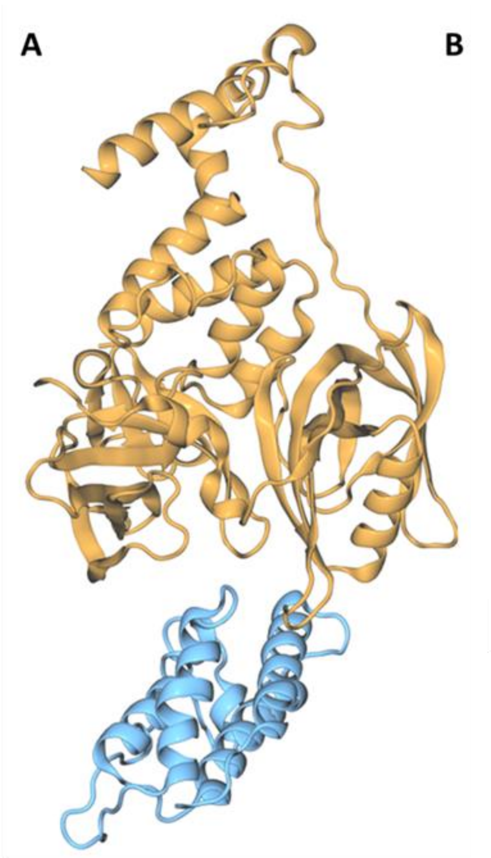


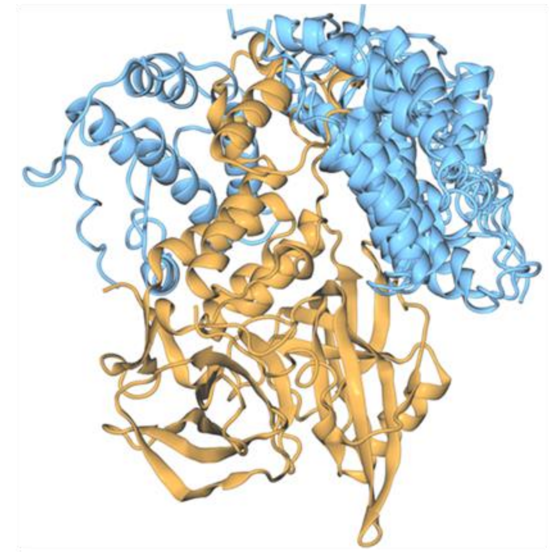


C

| Ranking | Docking Accuracy |
|---|---|
| Top1 | 0.00 |
| Top3 | 0.00 |
| Top5 | 0.08 |

**Figure S7** - Calculations using AlphaFold2-Multimer. A) Reference structure from 6BOY, where the PROTAC molecule dBET6 was removed for clarity, illustrating the interaction interface between cereblon (orange) and the BRD4 bromodomain (blue). B) All five models produced by AF2-Multimer, superimposed on cereblon. C) Table showing the docking accuracy calculated for all 13 systems using AF2-Multimer.
